# Characterization of regulatory transcriptional mechanisms in hepatocyte lipotoxicity

**DOI:** 10.1101/2021.03.24.436772

**Authors:** Joaquín Pérez-Schindler, Elyzabeth Vargas-Fernández, Bettina Karrer-Cardel, Danilo Ritz, Alexander Schmidt, Christoph Handschin

## Abstract

Non-alcoholic fatty liver disease is a continuum of disorders among which non-alcoholic steatohepatitis (NASH) is particularly associated with a negative prognosis. Hepatocyte lipotoxicity is one of the main pathogenic factors of liver fibrosis and NASH. However, the molecular mechanisms regulating this process are poorly understood. The main aim of this study was to dissect transcriptional mechanisms regulated by lipotoxicity in hepatocytes. We achieved this aim by combining transcriptomic, proteomic and chromatin accessibility analyses from human liver and mouse hepatocytes. This integrative approach revealed several transcription factor networks deregulated by NASH and lipotoxicity. To validate these predictions, genetic deletion of the transcription factors MAFK and TCF4 was performed, resulting in hepatocytes that were better protected against saturated fatty acid oversupply. MAFK- and TCF4-regulated gene expression profiles suggest a mitigating effect against cell stress, while promoting cell survival and growth. Moreover, in the context of lipotoxicity, some MAFK and TCF4 target genes were to the corresponding differentially regulated transcripts in human liver fibrosis. Collectively, our findings comprehensively profile the transcriptional response to lipotoxicity in hepatocytes, revealing new molecular insights and providing a valuable resource for future endeavours to tackle the molecular mechanisms of NASH.

## Introduction

Non-alcoholic fatty liver disease (NAFLD) is a spectrum of disorders that progresses from NAFL (hepatic steatosis) to non-alcoholic steatohepatitis (NASH), leading to cirrhosis and liver cancer (1). NASH is characterized by chronic liver damage, inflammation and fibrosis, a stage that significantly increases morbidity and mortality (1). Several genetic and environmental factors (e.g. a sedentary lifestyle) drive the development of NAFLD, among which the excessive supply of saturated fatty acids to the liver and, consequently, hepatocyte lipotoxicity is the main risk factor for NASH (1, 2). Importantly, hepatocyte lipotoxicity is the leading cause of liver fibrosis that remains as the main predictor of liver-related death (1, 2). However, therapies to target this process are scarce, as evidenced by the lack of approved medications for the treatment of NASH (3).

Although the molecular underpinnings controlling hepatocyte lipotoxicity are elusive, key biological processes such as oxidative stress, endoplasmic reticulum (ER) stress and inflammation are known to play a central role (1, 2). Interestingly, obesogenic diets extensively remodel the transcriptome, epigenome and chromatin accessibility landscape of mouse liver (4–7). These studies suggest that disruption of transcriptional networks and chromatin function are fundamental to pathogenic progression. Such transcriptional networks are regulated by transcription factors that, upon activation, are recruited to DNA regulatory elements located in regions of accessible chromatin (8). Despite the great therapeutic potential, the transcriptional mechanisms regulating hepatocyte lipotoxicity remain virtually unknown. Therefore, the main aim of this study was to characterize gene regulatory networks deregulated by saturated fatty acids oversupply in hepatocytes. We achieved this aim by implementing a multi-omics approach in which human and mouse datasets were integrated to uncover conserved transcriptional responses.

## Results

### NASH and lipotoxicity share a common molecular signature

To discover conserved hepatocyte-specific mechanisms linked to lipotoxicity and, potentially, NASH development, we implemented a multi-omics strategy in which we integrated analyses from human liver biopsies and mouse hepatocytes. Liver tissue was obtained from healthy control (CON) subjects and NASH patients suffering from obesity and diabetes (Figure 1A). In addition, lipotoxicity was investigated in FL83B mouse hepatocytes that were stimulated with bovine serum albumin (BSA) or BSA conjugated with the saturated fatty acid palmitate (PAL) for 24 h. As expected, PAL induced a dose-dependent increase in hepatocyte cytotoxicity (Figure 1B). We found that while a low dose of PAL increases the accumulation of intracellular lipids, incubating cells with a higher dose had no significant effect (Figure 1C). Consistently, a lower capacity to incorporate saturated fatty acids into triglyceride has been demonstrate to mediate the cytotoxic effect of PAL overload (9). Therefore, all of the following experiments were carried out under high PAL concentration (0.4 mM).

**Figure 1.**
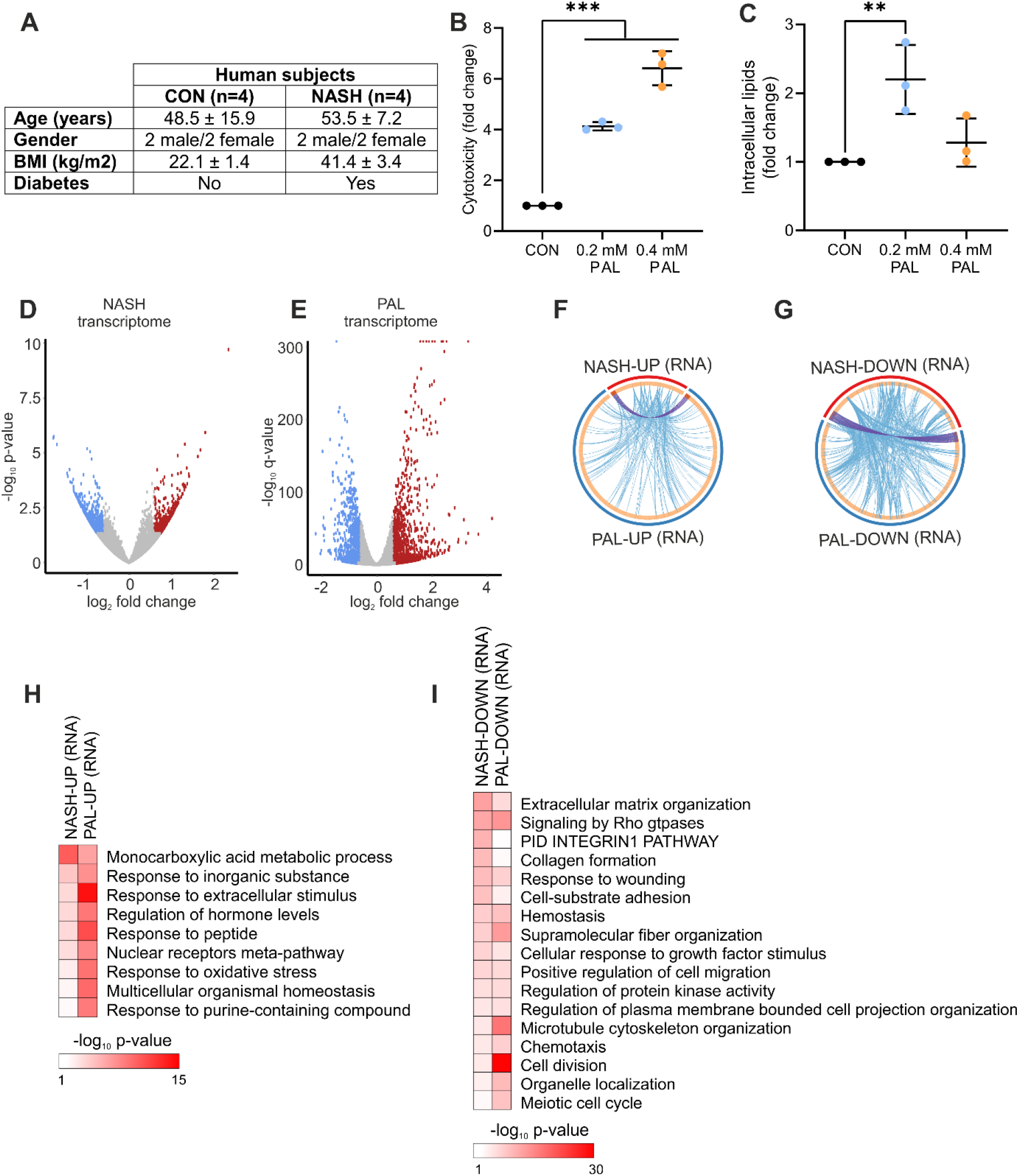
Shared transcriptional response in NASH and lipotoxicity. (A) Characteristics of human subjects. (B and C) Mouse hepatocytes were stimulated with BSA (control) or PAL for 24 h, and (B) cytotoxicity and (C) intracellular lipid content were measured (n=3 independent experiments; values are mean ± SD; **p < 0.01, ***p < 0.001). (D and E) Volcano plots showing differentially regulated genes in (D) human liver from NASH patients (n=4 per group) and (E) mouse hepatocytes stimulated with PAL for 24 h (n=3 per group). Blue and red dots denote significantly down- and up-regulated genes, respectively. (F and G) Circular plots showing the gene (purple lines) and functional (blue lines) overlap between (F) up- and (G) down-regulated genes in human NASH and mouse hepatocyte lipotoxicity. (H and I) Gene ontology analysis showing biological processes significantly regulated in both human NASH and mouse hepatocyte lipotoxicity (H) up- and (I) down-regulated genes.

We initially assessed whether human liver from NASH patients and mouse hepatocyte lipotoxicity share a common transcriptional response. In humans, NASH was associated with the up- and down-regulation of 422 and 615 genes, respectively (Figure 1D). Compared to these chronic changes in human NASH, mouse hepatocytes exhibited a stronger acute transcriptional response as evidenced by a higher number of differentially expressed genes (DEG), comprising 1744 and 1396 up- and down-regulated genes, respectively (Figure 1E). Although there was a small overlap of DEG between human NASH and hepatocyte PAL treatment (Figure 1F and 1G; purple lines), we observed a high degree of functional overlap reflected by a large number of genes belonging to the same ontology terms (Figure 1F and 1G; blue lines). Consistent with the induction of lipotoxicity and NASH, up-regulated genes were related to oxidative stress, nuclear receptor function and cellular response to stress (Figure 1H), while down-regulated genes were linked to processes such as the remodelling of the extracellular matrix, cell cycle progression and survival (Figure 1I).

We next assessed the liver and hepatocyte proteome to complement the transcriptome analysis and improve the comparison between NASH and lipotoxicity. We found that proteome remodelling was also milder in liver from NASH patients (up-regulated: 92 and down-regulated: 191) compared to the effects of PAL stimulation in mouse hepatocytes (up-regulated: 830 and down-regulated: 552) (Figure 2A and 2B). Similar to the transcriptome analysis, NASH and PAL proteomes showed a high level of functional overlap (Figure 2C and 2D; blue lines). A number of biological processes sensitive to both NASH and lipotoxicity were identified among up-regulated proteins, including oxidative stress and cellular response to stress, thus resembling the effects observed at the RNA level (Figure 2E). Moreover, our data demonstrate that down-regulated proteins were characterized by dysregulation of shared biological processes involved in protein folding, ER function, RNA processing and apoptosis (Figure 2F). Such functional overlap at the RNA and protein level suggest that lipotoxicity in mouse hepatocytes and in human NASH livers share a common molecular response.

**Figure 2.**
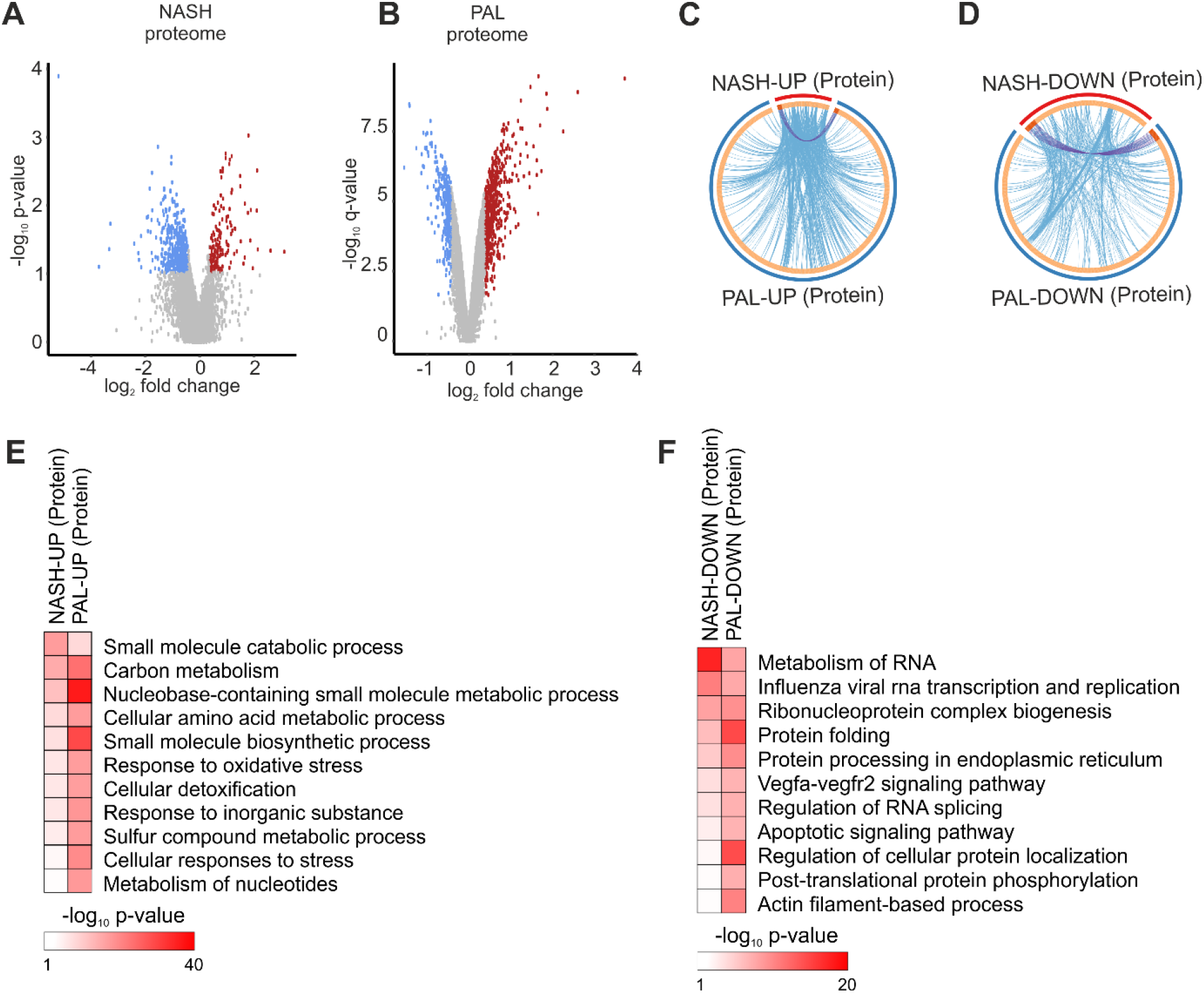
Shared proteome remodelling in NASH and lipotoxicity. (A and B) Volcano plots showing differentially regulated proteins in (A) human liver from NASH patients (n=4 per group) and (B) mouse hepatocytes stimulated with PAL for 24 h (n=3 per group). Blue and red dots denote significantly down- and up-regulated proteins, respectively. (C and D) Circular plots showing the protein (purple lines) and functional (blue lines) overlap between (C) up- and (D) down-regulated proteins in human NASH and mouse hepatocyte lipotoxicity. (E and F) Gene ontology analysis showing biological processes significantly regulated in both human NASH and mouse hepatocyte lipotoxicity (E) up- and (F) down-regulated proteins.

### NASH and lipotoxicity remodel the chromatin accessibility landscape

We next performed chromatin accessibility analysis by assay for transposase-accessible chromatin by sequencing (ATAC-seq) to elucidate transcriptional mechanisms mediating hepatocyte lipotoxicity and NASH. The total number of ATAC-seq peaks (representing regions of accessible chromatin) was similar between liver from CON and NASH (Figure 3A). In contrast, mouse hepatocytes treated with PAL had a lower number of peaks compared to the BSA control condition (Figure 3B). Both human and mouse ATAC-seq data showed that the vast majority of peaks were located in introns and DNA regulatory elements such as promoters and enhancers, at distal intergenic regions (Figure 3A and 3B). Interestingly, our data revealed that the proportion of genomic regions (e.g. introns and promoters) harbouring accessible 5 chromatin is to a large extent unaltered by NASH or lipotoxicity (Figure 3A and 3B). Globally, chromatin accessibility was slightly increased and strongly decreased in liver from NASH patients and in hepatocyte lipotoxicity, respectively (Figure 3C and 3D). Notably, in human livers, around 20% of the peaks were exclusively found in CON or NASH samples (Figure 3E and 3G). These data demonstrate that although the total number of ATAC-seq peaks is similar between groups, there are specific regions on the chromatin in which accessibility is lost (CON-specific peaks) or gained (NASH-specific peaks) during NASH development. In the context of mouse hepatocyte lipotoxicity, we observed a higher proportion of BSA-specific (35%) than PAL-specific (13%) peaks (Figure 3F and 3H), which is consistent with the global decrease in chromatin accessibility induced by PAL.

**Figure 3.**
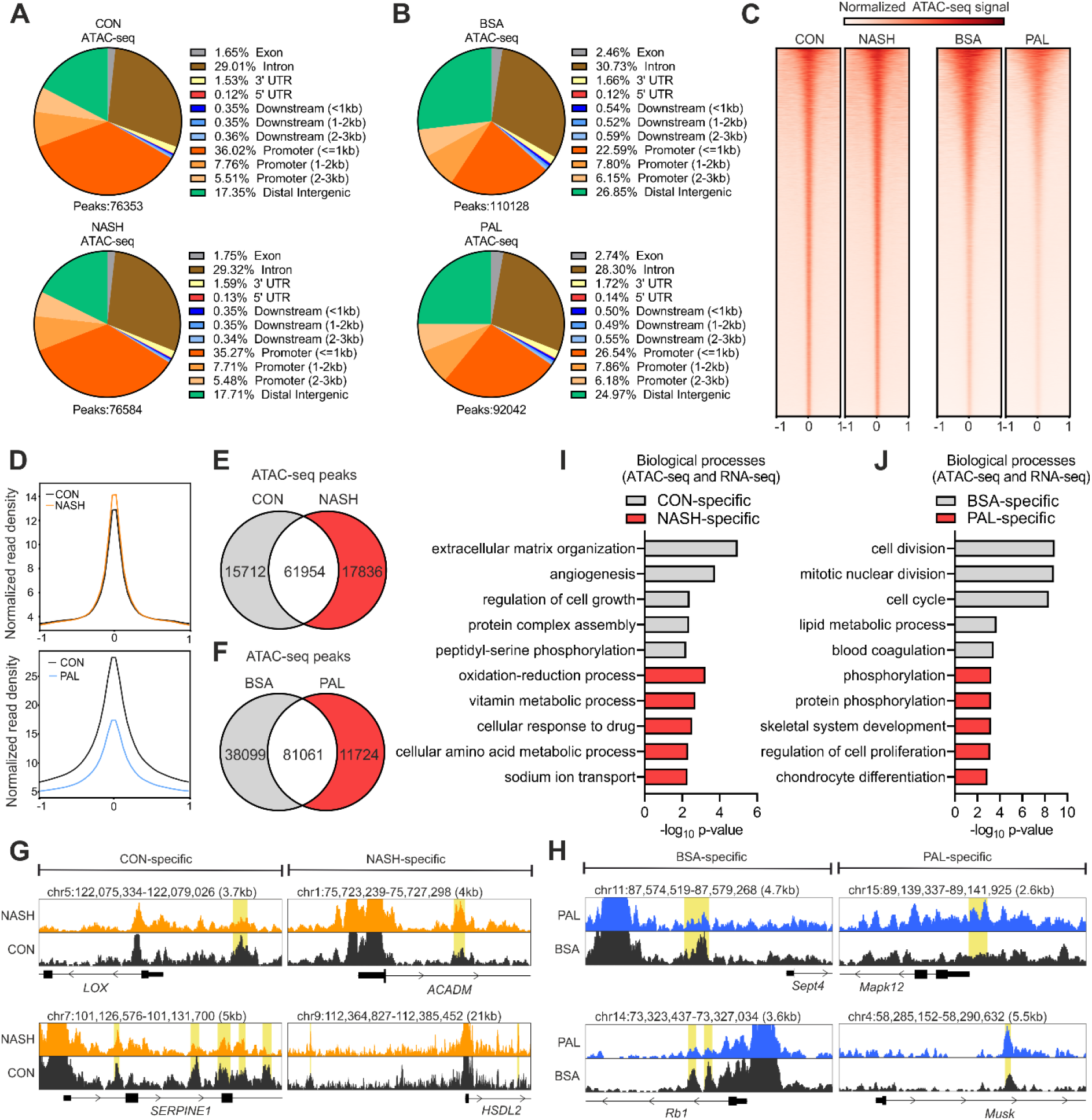
Chromatin accessibility is altered by NASH and lipotoxicity. (A and B) Annotation of ATAC-seq peaks in (A) human liver (n=4 per group) and (B) mouse hepatocytes (n=3 per group). (C) Heat maps of ATAC-seq peaks aligned to their centre ± 1 kb. (D) Density plots showing averaged normalized signal of ATAC-seq peaks aligned to their centre ± 1 kb. (E and F) Overlap of ATAC-seq peaks between (E) CON and NASH liver or (F) BSA and PAL hepatocytes. (G and H) Genome browser views of representative (G) CON-or NASH-specific peaks in human liver and (H) BSA-or PAL-specific peaks in mouse hepatocytes (marked yellow). (I and J) Gene ontology analysis showing biological processes regulated by genes linked to (I) CON-or NASH-specific peak in human liver and (J) BSA-or PAL-specific peaks in mouse hepatocytes.

To determine the link between changes in chromatin accessibility and gene expression, we identified down- and up-regulated genes associated with chromatin regions with decreased and increased accessibility, respectively. This analysis showed that repressed genes linked to CON-specific peaks were related to the regulation of cellular integrity, growth and vascularization, whereas up-regulated genes linked to NASH-specific peaks were mainly related to metabolic pathways (Figure 3I). Hepatocytes stimulated with PAL also exhibited a subset of down-regulated genes linked to BSA-specific peaks that we found to be primarily associated with the regulation of cell cycle and survival (Figure 3J). Moreover, genes up-regulated by PAL and linked to PAL-specific peaks were involved in the positive regulation of the unfolded protein response (UPR) and the development of fibrosis (Figure 3J), thus supporting the role of hepatocyte lipotoxicity in NASH development. Our data therefore further support the strong transcriptional component in both hepatocyte lipotoxicity and NASH.

### Lipotoxicity regulates a specific subset of transcription factors

Our findings strongly suggest that transcription factors play a central regulatory role in hepatocyte lipotoxicity, a process that holds great therapeutic potential for the treatment of NASH (3). We initially sought to identify such transcription factors via motif enrichment analysis of chromatin areas showing a loss (CON/BSA-specific) or gain (NASH/PAL-specific) in accessibility at distal intergenic regions (e.g. enhancers) and promoters. CON- and NASH-specific peaks had similar numbers of enriched transcription factor motifs at distal intergenic regions, but NASH was characterized by a strong enrichment of CTCF and CTCFL motifs (Figure 4A). In the context of hepatocyte lipotoxicity, we found that PAL treatment substantially decreased the number and enrichment of transcription factor motifs (CON-specific: 239 vs. PAL-specific: 55) at distal intergenic regions (Figure 4B). Analysis of promoters revealed similar results, where CON and NASH exhibited comparable number of enriched transcription factor motifs (Figure 4C), whereas PAL exerted a strong blunting effect (Figure 4D). We next defined the subset of transcription factor motifs shared between human liver and mouse hepatocytes at distal intergenic regions (CON/BSA: 103 and NASH/PAL: 25) and promoters (CON/BSA: 82 and NASH/PAL: 29). Interestingly, while most NASH/PAL transcription factor motifs were shared with CON/BSA, we found that 76% and 67% of transcription factor motifs were lost in distal intergenic regions and promoters, respectively, in the context of NASH and lipotoxicity (Figure 4E and 4F). In order to infer multi-protein transcriptional complexes dysregulated by NASH and lipotoxicity, we performed protein-protein interaction (PPI) network analysis. This approach revealed AP-1 transcription factor members as the only common significant PPI network in both distal intergenic regions and promoters (Figure 4G and 4H). In contrast, we found that CON/BSA-specific transcription factors formed distinct PPI networks (Figure 4G, 4H, S1A and S1B; distal intergenic: 5 and promoters: 4). Indeed, a number of transcription factors with known function in the regulation of liver function and NAFLD development were members of PPI networks at distal intergenic regions and promoters, including HNF4A, SMADs, ATF3, RXRA and ARNTL (Figure 4G, 4H, S1A and S1B) (10–13).

**Figure 4.**
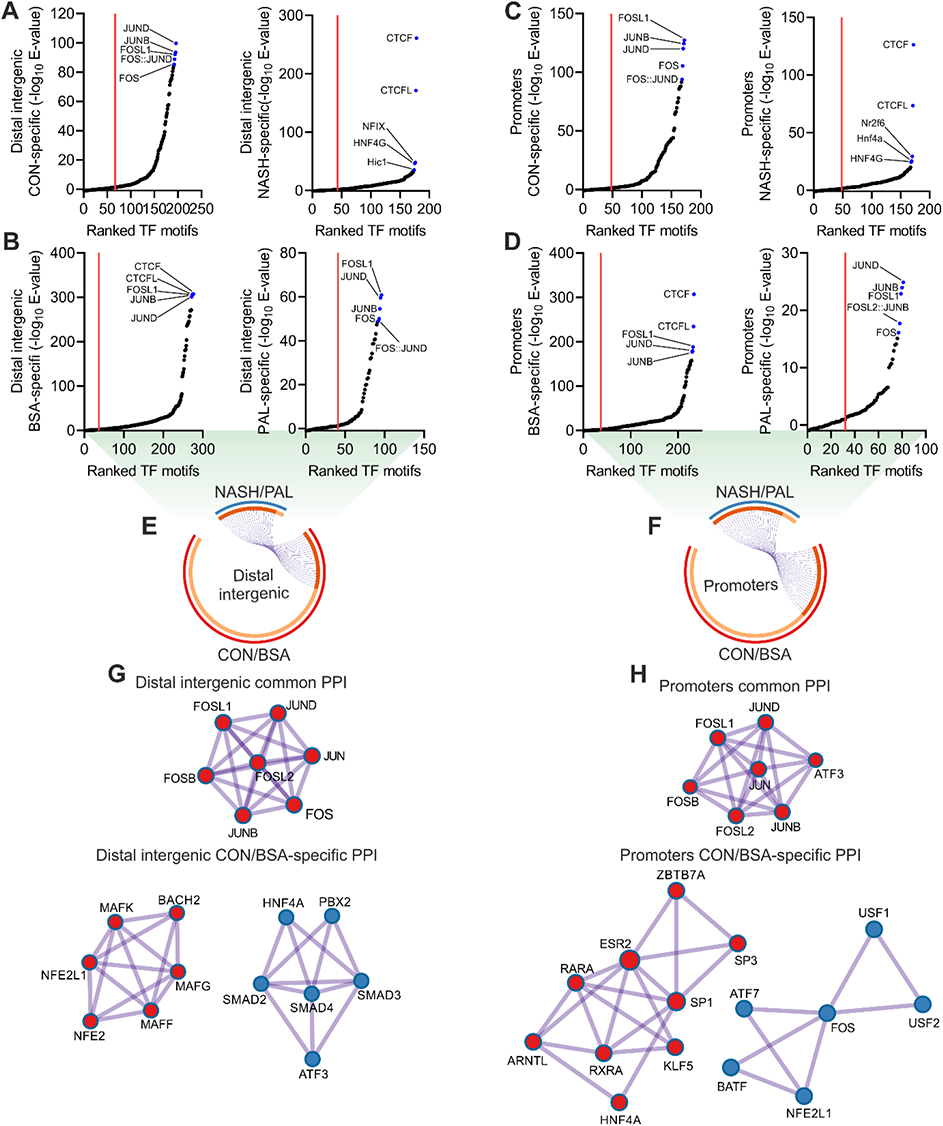
Conserved transcription factor networks at regulatory elements in NASH and lipotoxicity. (A-D) Transcription factor motif enrichment analysis in chromatin regions showing a loss (CON/BSA-specific) or gain (NASH/PAL-specific) in accessibility at (A and B) distal intergenic regions and (C and D) promoters in the context of NASH or PAL treatment (red line denotes significance cut-off of E-value < 0.05). (E and F). Circular plots showing conserved transcription factors (purple lines) shared between CON/BSA- and NASH/PAL-specific chromatin regions at (E) distal intergenic regions and (F) promoters. (G and H) Protein-protein interaction (PPI) network analysis of common and CON/BSA-specific transcription factors at (G) distal intergenic regions and (H) promoters.

To refine the identification of relevant transcription factor candidates in an unbiased manner, we performed transcription factor motif enrichment analysis with each of the independent datasets generated from human liver and mouse hepatocytes. The overlap between these datasets uncovered 227 motifs representing 170 transcription factors commonly enriched both in human liver and mouse hepatocytes (Figure 5A). We next used our transcriptomic data to perform Integrated System for Motif Activity Response Analysis (ISMARA), which specifically infers transcription factor activity at proximal promoter regions located 500 base pairs upstream and downstream of transcription start sites (14). This approach identified 36 and 172 motifs representing 63 and 249 transcription factors predicted to have altered activity in NASH and lipotoxicity, respectively. By integrating these analyses, we found 35 transcription factors deregulated by NASH and lipotoxicity, among which a subset of 13 also exhibited motif enrichment within regions of accessible chromatin (Figure 5A). Clustering analysis based on ISMARA-predicted activity showed a small fraction of transcription factors in which NASH and PAL induced the same effect (Figure 5B). Assessment of changes in the RNA levels of these transcription factors implies that, generally, the predicted activity does not directly relate to gene expression patterns (Figure 5B). Transcription factors sharing the same response might reflect redundant regulators due to similarities in their corresponding binding motifs on the DNA. We thus performed clustering analysis based on the consensus sequence of such motifs. Although we were able to define 5 distinct clusters, they did not match clustering patterns based on inferred transcription factor activity (Figure S2A). This suggests that most of these transcription factors regulate unique transcriptional networks related to hepatocyte lipotoxicity.

**Figure 5.**
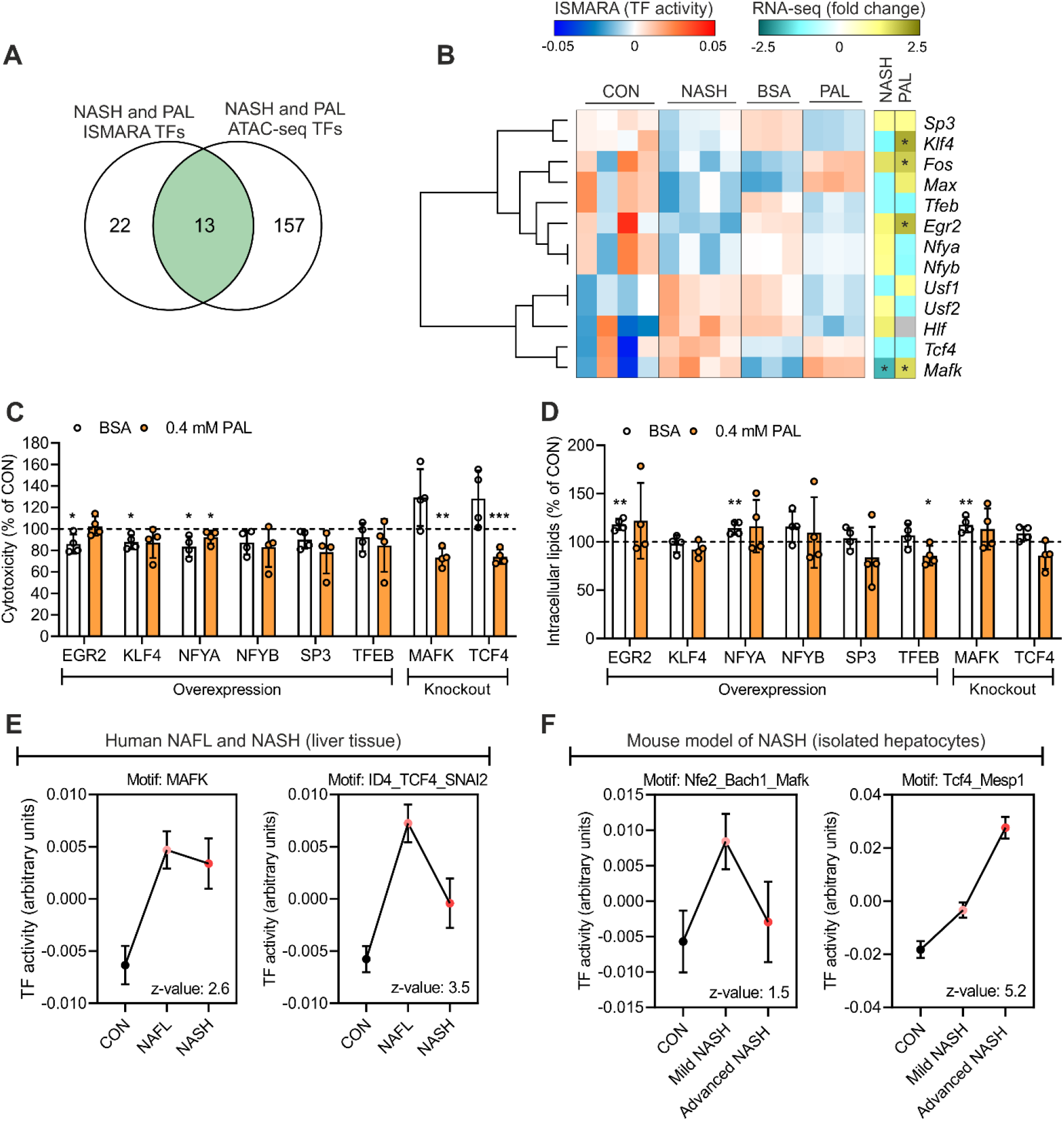
Omics data integration revealed candidate lipotoxicity-sensitive transcription factors at proximal promoters. (A) Overlap between all ISMARA (RNA-seq) and motif enrichment analysis (ATAC-seq) predicted transcription factors (TFs). (B) Heat map with ISMARA-predicted activity (left panel; clustering based of predicted TF activity) and gene expression changes (right panel; asterisk denotes a significant change) of TFs regulated in both systems. (C) Cytotoxicity and (D) intracellular lipid content measurement following BSA or PAL stimulation for 24 h in mouse hepatocytes with either overexpression (OE) or knockout (KO) of candidate TFs. Data are expressed as percentage of control cells (CON; corresponding to 100% denoted by the horizontal dashed line) undergoing the same treatment (n=4 independent experiments; values are mean ± SD; *p < 0.05, **p < 0.01 and ***p < 0.001). (E and F) ISMARA-predicted activity of MAFK and TCF4 in (E) human liver from CON subjects or NAFL and NASH patients (GEO accession: GSE126848), and (F) hepatocytes isolated from a mouse model of diet-induced NASH (GEO accession: GSE162876).

Subsequently, we selected a subset of transcription factor candidates that exhibit equal changes in activity both in NASH and lipotoxicity. We then either overexpressed (OE) repressed transcription factors (EGR2, KLF4, NFYA, NFYB, SP3 and TFEB) or knocked out (KO) those that were activated (MAFK and TCF4) in mouse hepatocytes (Figure S2B). This strategy allowed us to determine whether OE or KO of candidate transcription factors would confer protection against PAL-induced lipotoxicity. Only NFYA OE induced a minor decrease in the cytotoxic response to PAL (Figure 5C). We found that genetic deletion of either MAFK or TCF4 was sufficient to lower the cytotoxic response to PAL in mouse hepatocytes (Figure 5C). When TFEB was OE, we observed a slight decreased content of intracellular lipids (Figure 5D), whereas OE or KO of the other transcription factors had no effect on intracellular lipid storage (Figure 5D). Of note, to determine the reproducibility of our findings in the small sample size (n=4) of human NASH, we carried out an additional ISMARA analysis with a different RNA-seq dataset (GEO accession: GSE126848) from a large cohort of human liver samples, comprising a mild (NAFL, n=15) and advance (NASH, n=16) stage of NAFLD (15). Supporting our original results, this new transcription factor activity analysis showed that the predicted activity of MAFK and TCF4 was significantly increased in human liver from both NAFL and NASH patients (Figure 5E). Since the liver exhibits a high degree of cellular heterogeneity, we then performed ISMARA analysis using RNA-seq data from hepatocytes isolated from a diet-induced NASH mouse model (GEO accession: GSE162876) (16). Consistently, this analysis also predicts that NASH induces the activation of both MAFK and TCF4 specifically in hepatocytes (Figure 5F). Importantly, these independent validations of our findings further demonstrates that MAFK and TCF4 are lipotoxicity-sensitive transcription factors in hepatocytes, exhibiting a conserved response in liver from NAFLD patients.

### MAFK and TCF4 regulate distinct transcriptional networks linked to hepatocyte lipotoxicity

Next, we performed a proof of principle study to assess whether our data represent a useful resource to dissect mechanistic insights linked to the development of hepatocyte lipotoxicity and NASH. We achieved this by performing transcriptome analysis of CON and KO cells in the absence (BSA) or presence of PAL. TCF4 KO had a mild effect under basal condition, inducing the up- and down-regulation of 106 and 27 genes, respectively (Figure S3A). The total number of DEG induced by PAL was slightly higher in TCF4-KO cells, with about 26% of DEG being regulated in a TCF4-sensitive manner (Figure 6A, S3B and S3C). Moreover, direct comparison of cells stimulated with PAL revealed 80 and 97 gene showing a higher and lower expression in TCF4-KO cells, respectively (Figure S3D), representing commonly regulated genes also sensitive to TCF4 deletion. We performed gene ontology analysis with all DEG sensitive to PAL in CON and TCF4-KO cells to identify biological processes regulated in a TCF4-dependent manner. Interestingly, we observed that genetic deletion of TCF4 prevented induction of genes linked to cell death, stress and protein translation in response to PAL, while preventing the repression of genes promoting cell growth and survival (Figure 6B).

**Figure 6.**
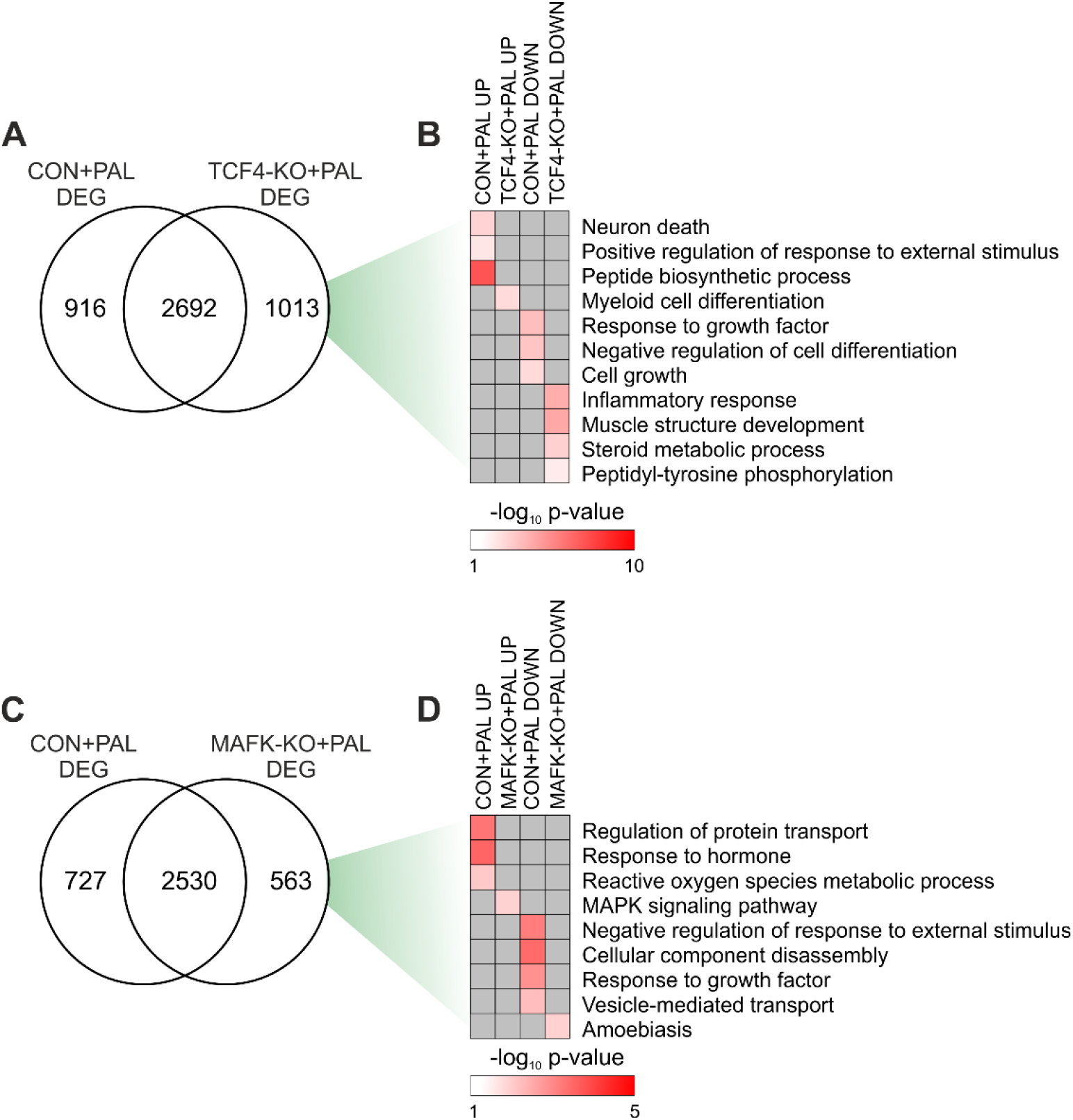
TCF4 and MAFK target genes are linked to hepatocyte lipotoxicity. (A and C) Overlap between DEG of CON and (A) TCF4-KO or (C) MAFK-KO hepatocytes stimulated with PAL for 24 h (n=3 per group). (B and D) Gene ontology terms exclusively regulated in CON and (B) TCF4-KO or (D) MAFK-KO cells.

We next carried out an analogous characterisation of MAFK-KO hepatocytes in the context of lipotoxicity. KO of MAFK under basal conditions resulted in the up- and down-regulation of 37 and 30 genes, respectively (Figure S3E). Transcriptome remodelling of PAL-stimulated cells was slightly higher in CON and, similarly to TCF4-KO cells, about 20% of DEG were regulated in a MAFK-sensitive manner (Figure 6C, S3F and S3G). We additionally found 21 and 17 up- and down-regulated genes, respectively, in MAFK-KO cells when compared to CON, both following PAL stimulation (Figure S3H). Similarly, the enrichment of terms associated to cell stress, toxicity and blunted growth was prevented in MAFK-KO cells (Figure 6D). Therefore, our data suggest that KO of both TCF4 and MAFK induces distinct gene expression signatures linked to the protective effect against lipotoxicity.

Considering the link between hepatocyte lipotoxicity and liver fibrosis in NASH prognosis, we subsequently defined the association between TCF4- and MAFK-sensitive genes and human NASH at different stages of fibrosis. We leveraged RNA-seq data (GEO accession: GSE135251) from a large human study comprising liver samples from NASH patients with moderate fibrosis (F2), severe fibrosis (F3) and cirrhosis (F4) (Figure 7A) (17). We found 90 and 16 up- and down-regulated genes, respectively, shared in humans NASH F4 and mouse hepatocyte lipotoxicity (Figure 7B). Among these genes, the up- and down-regulation of 17 and 3 was controlled by TCF4, respectively (Figure 7B). Interestingly, the expression of TCF4-sensitive genes appears to be directly related to liver fibrosis, where a higher degree of fibrosis is associated with a stronger effect in human liver (Figure 7C). While CON hepatocytes stimulated with PAL recapitulated the expression patterns of a subset of genes observed in liver from NASH F4 patients, genetic deletion of TCF4 was able to counteract such a response (Figure 7C). Similarly, when compared to human NASH F4, we identified 13 and 2 up- and down-regulated genes, respectively, regulated in a MAFK-depended manner (Figure 7D). MAFK-sensitive gene regulation was also linked to fibrosis in human liver, with genetic deletion of this transcription factor attenuating the effects of lipotoxicity (Figure 7E). These preliminary loss-of-function studies in mouse hepatocytes demonstrate that our results can be efficiently used for future data-integrated approaches to further dissect the molecular underpinnings of hepatocyte lipotoxicity and NASH.

**Figure 7.**
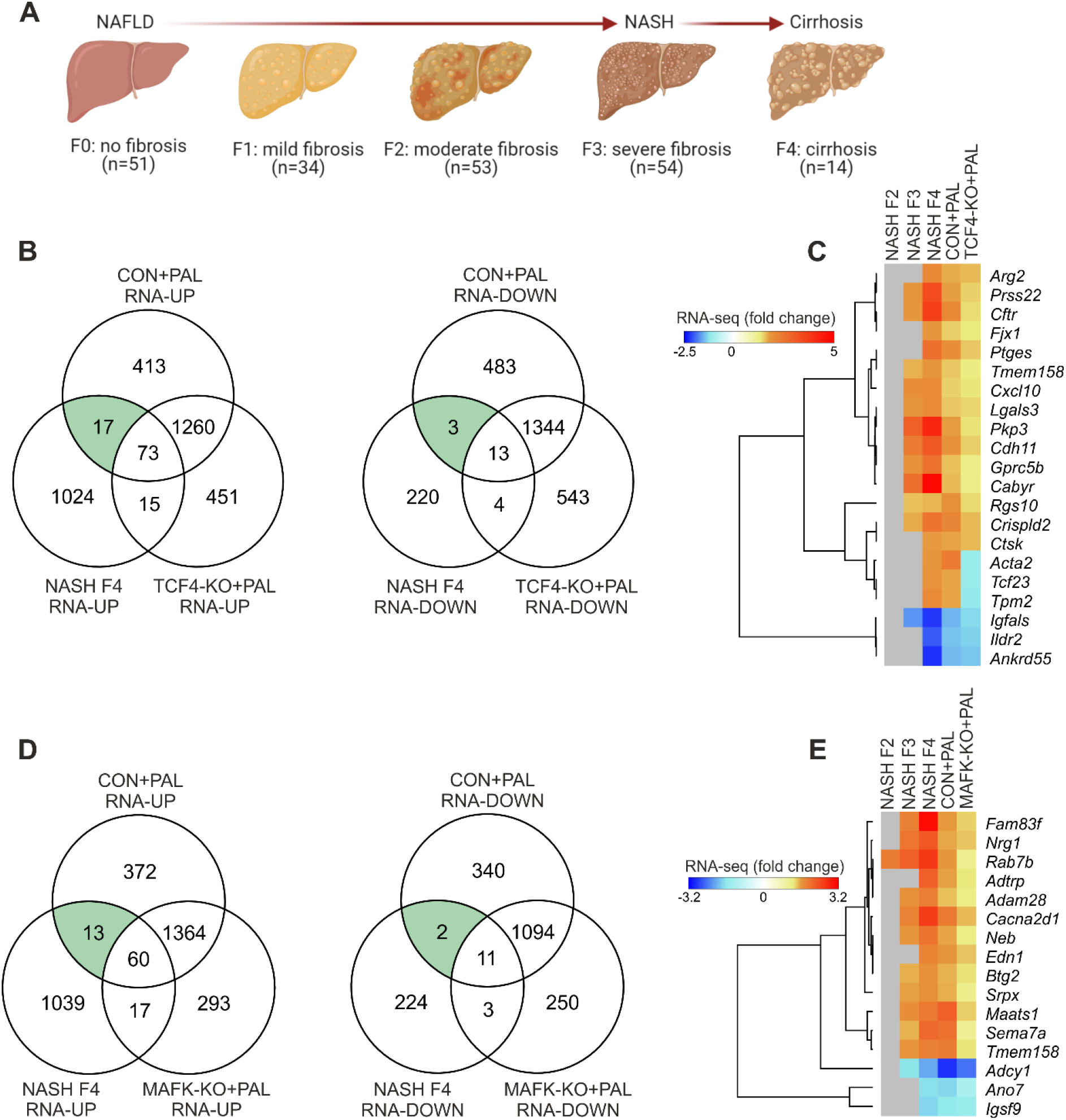
Overlap between TCF4 and MAFK target genes and human liver fibrosis. (A) Schematic representation of liver fibrosis stages from which samples used for downstream data integration were obtained (prepared with Biorender). (B and D) Overlap between DEG in human liver from NASH patients with cirrhosis (F4; GEO accession: GSE135251) and PAL-stimulate CON and (B) TCF4-KO or (D) MAFK-KO mouse hepatocytes. (C and E) Heat maps and clustering analysis of fold changes of (C) TCF4-or (E) MAFK-sensitive genes regulated both in human NASH F4 and mouse hepatocyte lipotoxicity, including data from human liver from NASH patients with moderate (F2) and severe (F3) fibrosis (grey denotes genes not significantly regulated in human NASH).

## Discussion

The development of hepatocyte lipotoxicity plays a crucial role in the development liver fibrosis associated to NASH, for which to date there are no approved medications (3). The overload of intracellular lipids in hepatocytes due to an oversupply of saturated fatty acids and/or enhanced de novo lipogenesis (e.g. high fructose consumption) induces the acute activation of pathway promoting cellular injury and death such as the UPR, oxidative stress and pro-inflammatory signals (1). However, the molecular underpinnings by which this process is regulated at the transcriptional level in hepatocyte has been poorly characterized, which to some extent has limited the identification of therapeutic targets for NASH. The current study provides a comprehensive profiling of the effects of lipotoxicity on the hepatocyte transcriptome, proteome and chromatin accessibility landscape, revealing several transcriptional networks with potential regulatory function conserved between mouse and human.

Beyond elucidating transcriptional mechanism in hepatocyte lipotoxicity, our findings can be leveraged to identify molecular insights with potential relevance in human disease. However, a key limitation of this study is the small number of human samples, for which we integrated our data with those from two and one independent human and mouse studies, respectively, comprising samples from liver biopsies from NAFL and NASH patients or mice. Thereby, a very stringent stratification of transcriptional programs and regulators was achieved. While this stringency might result in false negative calling of candidates due to the focus on hepatocytes and acute effects of PAL, this combined approach will reveal transcription factors featuring a regulation and function with potential relevance for the human liver biology and disease. Importantly, the integration of transcriptome and chromatin accessibility analyses allowed us to discover lipotoxicity-sensitive transcription factor networks and their downstream biological processes in hepatocytes.

Liver metabolism and function is regulated by a variety of transcriptional factors, some of which have been implicated in the development of NAFLD (10, 18). The pathogenic relevance of transcription factors is supported by several studies demonstrating that NAFLD is associated with altered chromatin function and, thus, gene expression (4–7). By performing a small pilot study, we identified TCF4 and MAFK as lipotoxicity-sensitive transcription factors in hepatocytes. To our knowledge, none of these transcription factors have been implicated in the development of hepatocyte lipotoxicity. However, a recent preprint article reports that liver from NASH-prone patients exhibit an enrichment of several transcription factor motifs located at enhancers that are shared with our data, including MAFK, TCF3 (with similar motif to TCF4), JUND, ATF3 and BACH2 among others (19). Of particular interest, we found that, in the context of lipotoxicity, TCF4 and MAFK regulated the expression of a subset of genes regulating cell stress, survival and homeostasis. For instance, TCF4-KO cells did not show an increase and decrease of genes linked to enhanced protein synthesis and cell growth, respectively. An abnormal increase in protein synthesis can overload the ER and thereby activate the UPR that consequently impairs cell growth and viability (20, 21). MAFK-sensitive genes did reveal a similar effects, characterized by the prevention of up- and down-regulation of genes involved in oxidative stress and cell growth, respectively. Therefore, our global unbiased analyses revealed a wide collection of MAFK- and TCF4-sensitive gene providing novel mechanistic insight into hepatocyte lipotoxicity.

An important strategy of our study was to determine candidate transcription factors and downstream target genes exhibiting a conserved response to NASH/lipotoxicity between human liver and mouse hepatocytes. Candidates showing such level of conservation are expected to play a central role in the regulation of hepatocyte function, with a possible link to disease development. In line with this idea, we found several genes with a known regulatory role in liver fibrosis (*Sema7a*, *Cdh11*) (22–24) and NAFLD development (*Adam28*, *Lgals3*, *Cxcl10*) (25–28) among MAFK- and TCF4-sensitive genes that are also regulated in human NASH. Notably, while the TCF4-sensitive gene *Cxcl10* has been proposed as a NASH biomarker (28), the *Lgals3* (also known as *Galectin* 3) inhibitor Belapectin is a promising medication for NASH currently under investigation and development for phase 3 clinical trials (3, 29). Collectively, this proof of principle study demonstrate that the datasets generated in this study are a valuable resource to dissect molecular mechanisms by which saturated fatty acids oversupply induces hepatocyte cytotoxicity, while uncovering genes with potential clinical relevance. In particular, we predict that integration of our data with work leveraging cell type-specific analyses (e.g. scRNA-seq) will be of particular relevance. However, future studies will be required to further investigate the relevance and therapeutic potential of the transcriptional networks identified in this study, for instance using preclinical mouse models of NASH.

In conclusion, our findings have uncovered a new layer of complexity in hepatocyte lipotoxicity, where transcriptional networks sensitive to saturated fatty acids promote cell death. By performing thorough comparisons to human datasets and experimental validation, we identified a number of transcriptional networks and biological processes associated with the development of hepatocyte lipotoxicity. Furthermore, our data provide a rich resource to generate new hypotheses for future studies aiming at tackling liver lipotoxicity, the central pathogenic factor of NASH and liver-related death.

## Materials and methods

### Ethics statement

Human liver tissue was collected with appropriate consent and ethical approval obtained by Sekisui XenoTech.

### Human samples

Fresh frozen and pulverized human liver samples were purchased from Sekisui XenoTech. Samples were obtained from normal (lot #: H1281, H1296, H1310 and H1336) and non-alcoholic steatohepatitis (lot #: H0847, H0958, H1027 and H1060) donors.

### Cell culture and palmitate treatment

FL83B mouse hepatocytes (ATCC^®^, #CRL-2390™) were grown in F-12K medium (Thermo Fisher Scientific, #21127030) supplemented with 10% fetal bovine serum (growth medium). Cells were maintained at 37°C, 95% O_2_ and 5% CO_2_.

A stock solution of 25 mM sodium palmitate (PAL) (Sigma-Aldrich, #P9767) was prepared in serum-free F-12K media containing 1% fatty acid free bovine serum albumin (BSA) (Sigma-Aldrich, #A8806), which was incubated at 70°C for 30 min shaking at 1000 rpm before use. Next, cells were incubated in 0, 200 and 400 µM PAL diluted in growth medium for 24 h. One percent BSA diluted in growth medium was used as control.

### Cytotoxicity assay

Cytotoxicity was measured with CellTox™ Green Cytotoxicity Assay (Promega, #G8742), according to the manufacturer’s instructions.

### Intracellular lipid staining

Cells were incubated with a staining solution containing 1 μg/ml of Hoechst 33342 (Thermo Fisher Scientific, #H3570) and 500 ng/ml of Nile Red (Sigma-Aldrich, #N3013) in phosphate buffered saline (PBS) for 20 min covered from light at 37 °C and 5 % CO_2_. Next, staining solution was removed and cells were washed twice with PBS. Fresh PBS was added before measuring Nile Red (excitation/emission = 488/550 nm) and Hoechst 33342 (excitation/emission = 350/461 nm) fluorescence with a microplate reader. Nile Red signal (lipids) was normalized to Hoechst 33342 signal (DNA) to account for differences in cell number.

### Transient transfections

Transfection of FL83B cells was performed using Opti-MEM™ (Thermo Fisher Scientific, #31985070) and polyethylenimine (Polysciences, # 23966). Plasmids and polyethylenimine were diluted in Opti-MEM™, following which they were mixed in a 1:3 ratio of ug DNA:ug polyethylenimine and incubated for 20 min at room temperature before adding to the cells. Cells were transfected 24 h after seeding with 0.1 or 1 μg per well of a 96 and 12 well plate, respectively, of pcDNA 3.1, pAd-Klf4 (Addgene, # 19770, gift from Konrad Hochedlinger), pFLAG/HA/mSp3 (VectorBuilder, #VB200204-1104aty), pFLAG/HA/mTfeb (VectorBuilder, #VB200204-1105cfn), pFLAG/HA/mEgr2 (VectorBuilder, #VB200204-1106upe), pFLAG/HA/mNfya (VectorBuilder, #VB200204-1107xne) or pFLAG/HA/mNfyb (VectorBuilder, #VB200204-1108efg) for a total of 48 h. Palmitate treatment was started 24 h after transfection.

### Generation of knockout cells

Gene knockout (KO) was achieved by using the CRISPR-Cas9 system. To KO the mouse Klf4 gene, FL83B cells were transfected as described above with the following plasmids: p-hCas9-mTcf4-gRNA#32572-mTcf4-gRNA#30405 (VectorBuilder, #VB200206-1079fgw) and p-hCas9-Scramble-gRNA1-Scramble-gRNA2 (VectorBuilder, #VB200206-1080bvj) as non-targeting control. Deletion of the mouse Mafk gene was performed using the Edit-R gene editing system (Horizon Discovery). First, stable Cas9 expression was induced by transducing FL83B cells with lentiCas9-Blast (Addgene, #52962-LV, gift from Feng Zhang) lentivirus at a multiplicity of infection (MOI) of 0.5 in growth medium containing 10 µg/ml of polybrene (Sigma-Aldrich, #107689). Next, cells were selected with 6 μg/ml of blasticidin (Thermo Fisher Scientific, #A1113903) and a monoclonal cell line was obtained via limiting dilution. Cas9 expressing FL83B cells were subsequently transduced with Mafk (#VSGM10144-246720110) or non-targeting control (#VSGC10215) lentiviral sgRNA at an MOI of 0.5 in growth medium containing 10 µg/ml of polybrene. Seventy two hours after Tcf4 or Mafk targeting, cells were selected with 1 μg/ml of puromycin (Thermo Fisher Scientific, #A1113803) to generate a polyclonal population of KO cells for downstream experiments.

### RNA purification and quantitative PCR (qPCR)

Liver tissue and cells were lysed in 1 ml of TRI Reagent (Sigma #T9424) and incubated for 5 min at room temperature. Aqueous phase was obtained with chloroform following the manufacturer’s instructions, following which RNA was purified and reverse transcribed using Direct-zol™ RNA MiniPrep (Zymo Research, #R2050) and iScript™ cDNA Synthesis Kit (Bio-Rad, #1708891), respectively. Relative changes in mRNA content was quantified by qPCR on a StepOnePlus system (Applied Biosystems) using Fast SYBR™ Green Master Mix (Thermo Fisher Scientific, #4385612). The ΔΔCT method was used for analysis, with TATA binding protein (*Tbp*) as endogenous control.

### RNA sequencing (RNA-seq)

Libraries from human liver tissue and wild type FL83B cells were prepared with TruSeq Stranded Total RNA Library Prep Gold (Illumina, #20020599), single-read sequencing was performed using the NextSeq 500 (Illumina). Libraries from knockout FL83B cells and their corresponding non-targeting controls were prepared with TruSeq Stranded mRNA Library Kit (Illumina, #20020595), pair-end sequencing was performed using the NovaSeq 6000 (Illumina). All data was analysed on the Galaxy platform (https://usegalaxy.eu/). Reads were trimmed with Trim Galore! (Galaxy version 0.4.3.1) and quality was assessed using FastQC (Galaxy version 0.72+galaxy1). Reads were aligned to the hg38 or mm10 version of the human and mouse genome, respectively, using STAR (Galaxy version 2.7.7a). Strand specificity and read counting was performed with Infer Experiment (Galaxy version 2.6.4.1) and featureCounts (Galaxy version 2.0.1), respectively. Next, we used DESeq2 (Galaxy version 2.11.40.6+galaxy1) for differential expression analysis (fold change ≥ 1.5, p-value < 0.05 for human liver tissue and q-value < 0.05 for FL83B cells) and the resulting data was annotated with Annotate DESeq2/DEXSeq output tables (Galaxy version 1.1.0). Overlap between different datasets was determined with Venny (version 2.1, https://bioinfogp.cnb.csic.es/tools/venny/) and volcano plots generated with Volcano Plot (Galaxy Version 0.0.3). Generation of circular plots and Gene Ontology (GO) analysis was performed with Metascape (https://metascape.org/gp/index.html#/main/step1) (30). Transcription factors activity analysis was achieved with ISMARA (https://ismara.unibas.ch/mara/), where z-value ≥ 1.5 was consider as significant. We used DESeq2 normalized counts to generate heat maps and for hierarchical clustering using Morpheus (https://clue.io/morpheus).

### Mass spectrometry analysis of whole cell proteome

Fifteen milligrams of powdered human liver tissue were lysed in 200 uL of lysis buffer (8M Urea, 50 mM Tris-HCl pH7.5, 150 mM NaCl and 1X Halt™ Protease Inhibitor Cocktail (Thermo Fisher Scientific, #87786)), vortexed at max speed for 15 sec and incubated for 30 min at 4°C with shaking at 14000 rpm. Next, samples were sonicated (amplitude 100%, Cycle 0.5) four times for 30 sec in a vial tweeter sonicator with 2 min pause on ice between cycles. Samples were then centrifuged for 13000 g for 10 min at 4°C, supernatant was transferred to a new tube and the pellet was homogenized in 50 μl of lysis buffer. Following an additional centrifugation at 13000 g for 10 min at 4°C, supernatant was combined with the previous supernatant. Samples were centrifuged a final time at 13000 g for 10 min at 4°C and supernatant was used for mass spectrometry. On the other hand, mouse hepatocytes were seeded in 6 well plates and harvested in 80 μl of lysis buffer per well (1% sodium deoxycholate (SDC), 0.1 M TRIS, 10 mM TCEP, pH = 8.5), following lysis with 10 cycles of sonication (Bioruptor, Diagnode). All samples were reduced for 10 min at 95°C and alkylated at 15 mM chloroacetamide for 30 min at 37°C. Proteins were digested by incubation with sequencing-grade modified trypsin (1/50 w/w; Promega, V5113) for 12 h at 37°C. Tryptic digests were acidified (pH<3) using TFA and cleaned up using iST cartridges (PreOmics, P.O.00027) according to the manufacturer’s instructions. Samples were dried under vacuum and stored at −20 °C.

Sample aliquots comprising 25 μg of peptides were labelled with isobaric tandem mass tags (TMT 10-plex, Thermo Fisher Scientific, #90110) as described previously (31). Shortly, peptides were re-suspended in 20 μl labelling buffer (2 M urea, 0.2 M HEPES, pH 8.3) and 5 μL of each TMT reagent were added to the individual peptide samples followed by a 1 h incubation at 25°C, shaking at 500 rpm. To quench the labelling reaction, 1.5 μL aqueous 1.5 M hydroxylamine solution was added and samples were incubated for another 10 min at 25°C shaking at 500 rpm followed by pooling of all samples. The pH of the sample pool was increased to 11.9 by adding 1 M phosphate buffer (pH 12) and incubated for 20 min at 25°C shaking at 500 rpm to remove TMT labels linked to peptide hydroxyl groups. Subsequently, the reaction was stopped by adding 2 M hydrochloric acid until a pH < 2 was reached. Finally, peptide samples were further acidified using 5 % TFA, desalted using Sep-Pak Vac 1cc (50 mg) C18 cartridges (Waters, #WAT054960) according to the manufacturer’s instructions and dried under vacuum.

TMT-labelled peptides were fractionated by high-pH reversed phase separation using a XBridge Peptide BEH C18 column (3,5 µm, 130 Å, 1 mm x 150 mm; Waters, #186003562) on an Agilent 1260 Infinity HPLC system. Peptides were loaded on column in buffer A (20 mM ammonium formate in water, pH 10) and eluted using a two-step linear gradient from 2% to 10% in 5 min and then to 50% buffer B (20 mM ammonium formate in 90% acetonitrile, pH 10) over 55 min at a flow rate of 42 µl/min. Elution of peptides was monitored with a UV detector (215 nm, 254 nm) and a total of 36 fractions were collected, pooled into 12 fractions using a post-concatenation strategy as previously described (32) and dried under vacuum.

Dried peptides were re-suspended in 0.1% aqueous formic acid and subjected to LC–MS/MS analysis using a Q Exactive HF Mass Spectrometer fitted with an EASY-nLC 1000 (Thermo Fisher Scientific) and a custom-made column heater set to 60°C. Peptides were resolved using a RP-HPLC column (75μm × 30cm) packed in-house with C18 resin (ReproSil-Pur C18–AQ, 1.9 μm resin; Dr. Maisch, r119.aq.) at a flow rate of 0.2 μLmin-1. The following gradient was used for separation of murine peptides: from 5% B to 15% B over 10 min to 30% B over 60 min to 45 % B over 20 min to 95% B over 2 min followed by 18 min at 95% B, whereas the following gradient was used for separation of human peptides: from 5% B to 15% B over 14 min to 30% B over 80 min to 45 % B over 26 min to 95% B over 2 min followed by 18 min at 95% B. Buffer A was 0.1% formic acid in water and buffer B was 80% acetonitrile, 0.1% formic acid in water.

The mass spectrometer was operated in DDA mode with a total cycle time of approximately 1 s. Each MS1 scan was followed by high-collision-dissociation (HCD) of the 10 most abundant precursor ions with dynamic exclusion set to 30 s. For MS1, 3e6 ions were accumulated in the Orbitrap over a maximum time of 100 ms and scanned at a resolution of 120,000 FWHM (at 200 m/z). MS2 scans were acquired at a target setting of 1e5 ions, maximum accumulation time of 100 ms and a resolution of 30,000 FWHM (at 200 m/z). Singly charged ions and ions with unassigned charge state were excluded from triggering MS2 events. The normalized collision energy was set to 35%, the mass isolation window was set to 1.1 m/z and one microscan was acquired for each spectrum.

The acquired raw-files were converted to the mascot generic file (mgf) format using the msconvert tool (part of ProteoWizard, version 3.0.4624 (2013-6-3)) and searched using MASCOT either against a murine database (consisting of 49434 forward and reverse protein sequences downloaded from Uniprot on 20141124) or a human database (consisting of 40832 forward and reverse protein sequences downloaded from Uniprot on 20181213) and 390 commonly observed contaminants. The precursor ion tolerance was set to 10 ppm and fragment ion tolerance was set to 0.02 Da. The search criteria were set as follows: full tryptic specificity was required (cleavage after lysine or arginine residues unless followed by proline), 3 missed cleavages were allowed, carbamidomethylation (C) and TMT6plex (K and peptide N-terminus) were set as fixed modification and oxidation (M) as a variable modification. Next, the database search results were imported into the Scaffold Q+ software (version 4.3.2, Proteome Software Inc.) and the protein false discovery rate was set to 1% based on the number of decoy hits. Proteins that contained similar peptides and could not be differentiated based on MS/MS analysis alone were grouped to satisfy the principles of parsimony. Proteins sharing significant peptide evidence were grouped into clusters. Acquired reporter ion intensities in the experiments were employed for automated quantification and statistical analysis using a modified version of our in-house developed SafeQuant R script v2.3 (31). This analysis included adjustment of reporter ion intensities, global data normalization by equalizing the total reporter ion intensity across all channels, summation of reporter ion intensities per protein and channel, calculation of protein abundance ratios and testing for differential abundance using empirical Bayes moderated t-statistics. The calculated p-values were corrected for multiple testing using the Benjamini−Hochberg method, with significance defined as fold change ≥ 1.3, p-value < 0.05 for human liver tissue and q-value < 0.05 for FL83B cells. Volcano plots were generated on the Galaxy platform (https://usegalaxy.eu/) with Volcano Plot (Galaxy Version 0.0.3). Generation of circular plots and Gene Ontology (GO) analysis was performed with Metascape (https://metascape.org/gp/index.html#/main/step1) (30). Overlap between different datasets was determined with Venny (version 2.1, https://bioinfogp.cnb.csic.es/tools/venny/).

### Assay for transposase-accessible chromatin by sequencing (ATAC-seq)

Nuclei from human liver (15 mg) were isolated using with the Nuclei EZ Prep kit (Sigma-Aldrich, #NUC101) and a glass dounce homogenizer (25 strokes with lose pastel and 25 strokes with tight pastel in ice-cold Nuclei EZ Lysis Buffer). Nuclei were filter through a 40 μm cell strainer and counted before ATAC-seq. On the other hand, Fl83B mouse hepatocytes were seeded in 60 mm plates 24 h before BSA or PAL treatment. Nuclei from human liver and mouse hepatocytes were then used for ATAC-seq and library preparation as previously described (33). Pair-end sequencing of the libraries was performed using the Illumina NextSeq 500. All data was analysed on the Galaxy platform (https://usegalaxy.eu/). Reads were trimmed with Cutadapt (Galaxy version: 1.16.5) and quality was assessed using FastQC (Galaxy version 0.72+galaxy1). Reads were aligned to the hg38 or mm10 version of the human and mouse genome, respectively, using Bowtie2 (Galaxy version 2.3.4.3+galaxy0). Low quality reads (phred < 30) were filtered out with Filter (Galaxy Version 2.4.1) and duplicated reads were removed with MarkDuplicates (Galaxy version 2.18.2.2). Next, we used Genrich (Galaxy version 0.5+galaxy2) for peak calling (q-value < 0.05), while ChIPseeker (Galaxy version, #1.18.0+galaxy1) was used to annotate peaks. The overlap between different ATAC-seq datasets was performed with bedtools Intersect intervals (Galaxy version 2.29.0). Heat map and density plots of ATAC-seq peaks were created with plotHeatmap (Galaxy version 3.0.2.0) and plotProfile (Galaxy version 3.1.2.0.0), respectively. Data was visualized on the Integrated Genome Browser-9.1.4 (34) to generate representative genome browser figures, for which BAM files were merged with Merge BAM Files (Galaxy version 1.2.0) and then normalized using bamCoverage (Galaxy version 3.0.2.0). CentriMo (version 5.1.0, https://meme-suite.org/meme/doc/centrimo.html) was used to perform motif enrichment analysis (E-value threshold < 0.05) (35). Generation of circular plots and protein-protein interaction network analysis was performed with Metascape (https://metascape.org/gp/index.html#/main/step1) (30). GO analysis was performed with DAVID 6.8 (https://david.ncifcrf.gov/). Finally, clustering of transcription factor motifs based in their consensus sequence was performed using STAMP (http://www.benoslab.pitt.edu/stamp/) (36).

### Statistics

All qPCR, cytotoxicity and intracellular lipid assays were performed at least three independent times each in triplicate. Number of replicates per experiment is indicated in the figure legend when appropriate. Values are expressed as mean ± SD. Statistical significance was determined with unpaired two-tailed t-tests, with significance considered with a p < 0.05. RNA-seq, ATAC-seq and mass spectrometry experiments were performed once with three to four biological replicates as indicated in figure legends. Statistical analysis of these experiments is described above in their corresponding sections.

### Data availability

RNA-seq and ATAC-seq data will be available at the Gene Expression Omnibus (GEO). Whole proteome analysis will be available at the Proteomics Identification Database (PRIDE). All other materials generated in this study are available from the corresponding authors upon request.

## Acknowledgements

We thank Christian Beisel (Genomics Facility Basel, ETH Zürich) and Philippe Demougin (Life Sciences Training Facility, Biozentrum, University of Basel) for technical help. We thank Eva Dazert for advice about processing of human liver samples (Biozentrum, University of Basel).

## Author contributions

JPS and CH conceived, designed and supervised the study. JPS, EVF, BKC, DR and AS performed experiments. JPS, DR, AS and CH performed data analysis and interpretation. JPS and CH wrote the manuscript. All authors reviewed the manuscript.

## Supplementary figures

**Figure S1.**
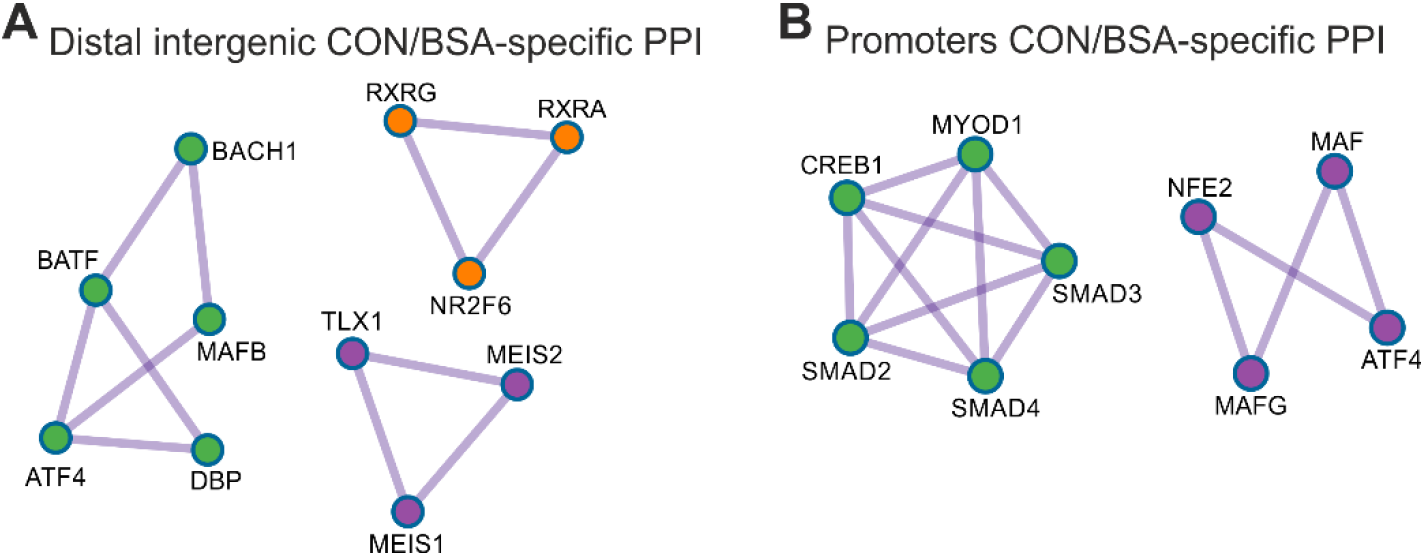
Protein-protein interaction (PPI) network analysis. (A and B) PPI networks of CON/BSA-specific transcription factors at (A) distal intergenic regions and (B) promoters.

**Figure S2.**
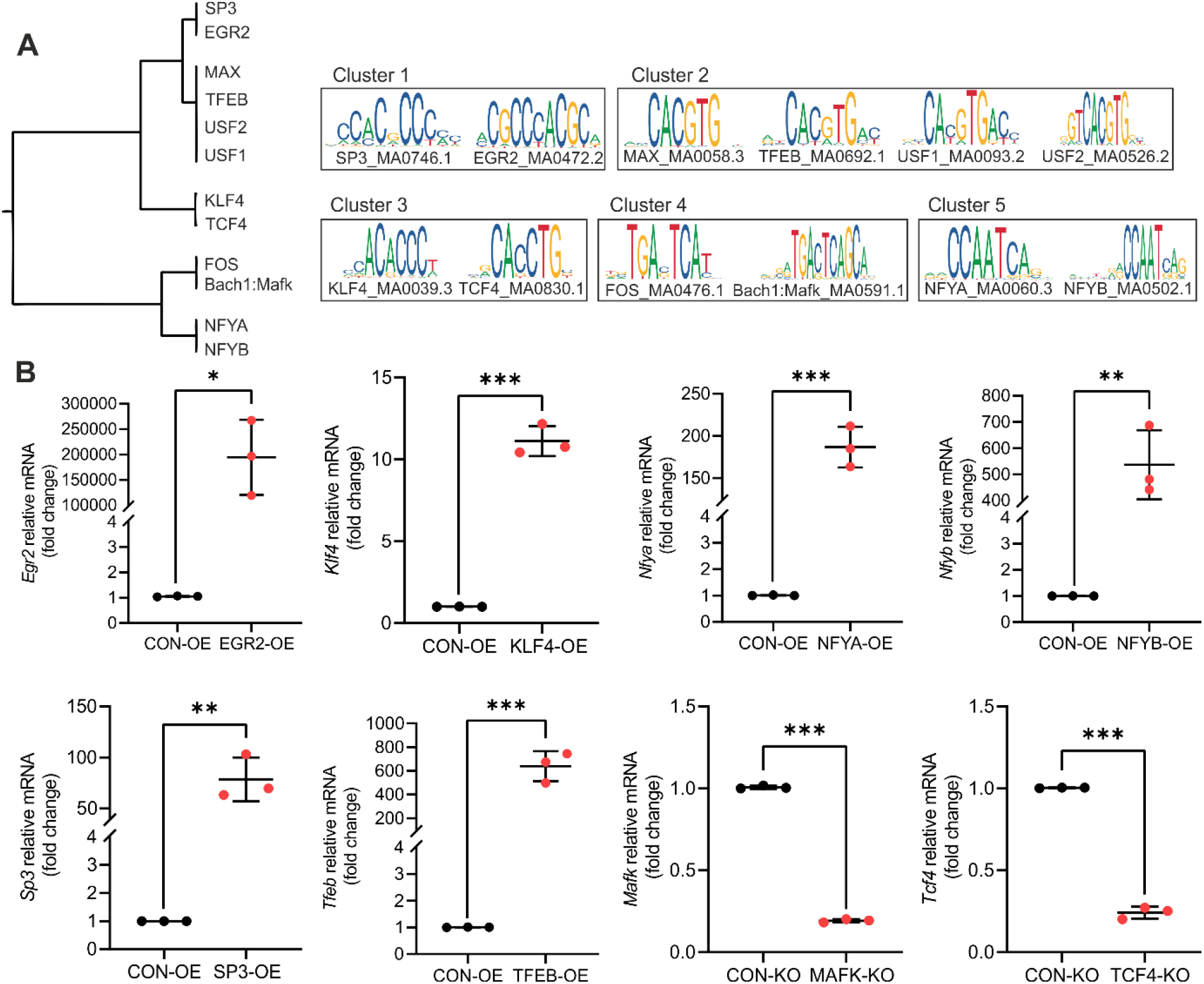
Lipotoxicity-sensitive transcription factors. (A) Transcription factor clustering analysis based on motif consensus sequence (left panel), with the motif logos comprised in the different clusters (right panel). (B) Transcript level of candidate transcription factors following overexpression (OE) or knockout (KO) in mouse hepatocytes compared to their corresponding controls (CON; n=3 independent experiments; values are mean ± SD; *p < 0.05, **p < 0.01 and ***p < 0.001)..

**Figure S3.**
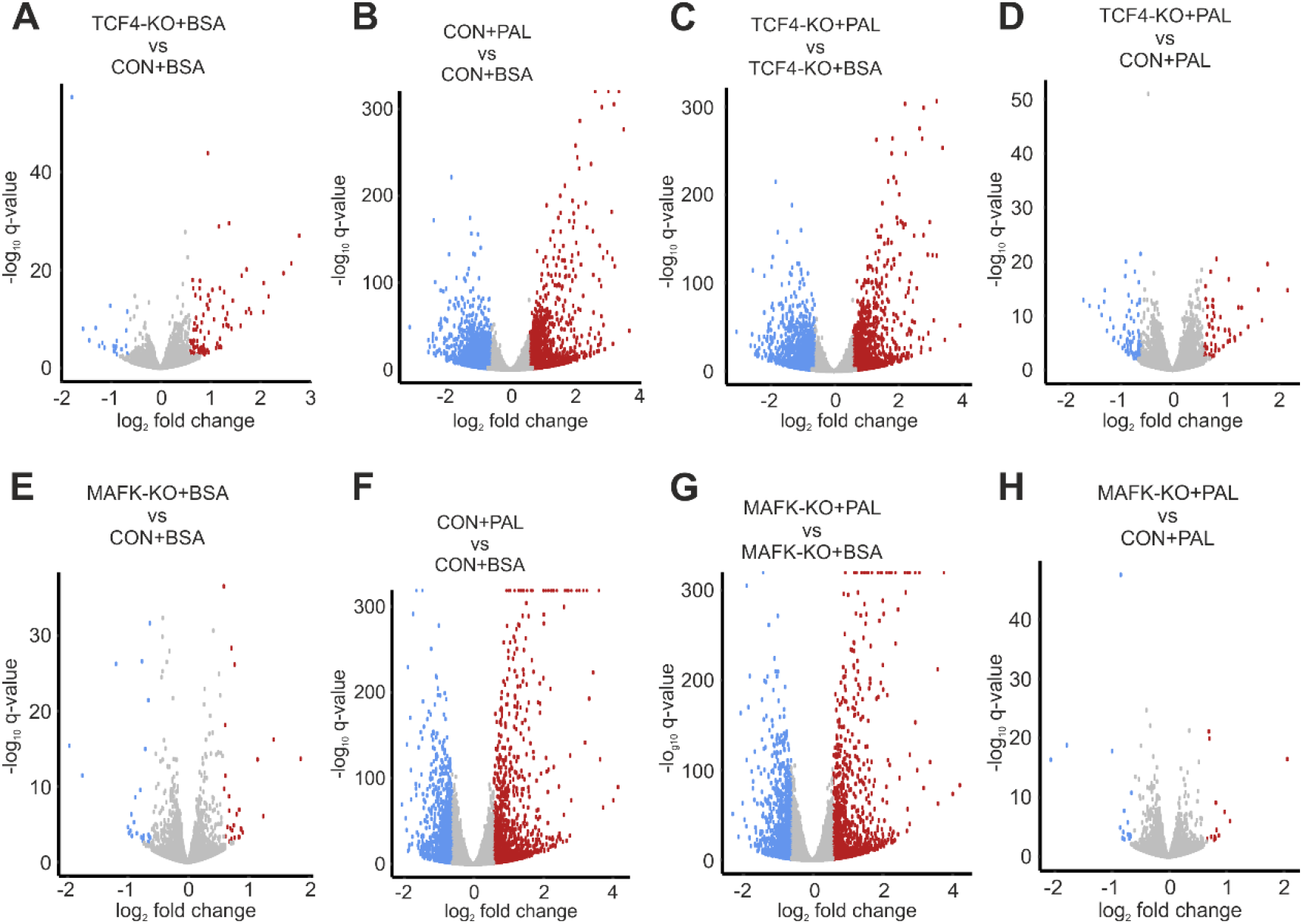
Transcriptomic analysis of MAFK and TCF4 knockout cells. (A-H) Volcano plots showing DEG under basal conditions and following PAL stimulation for 24 h in (A-D) TCF4-KO or (E-H) MAFK-KO mouse hepatocytes (n=3 per group; CON: control; blue and red dots denote significantly down- and up-regulated genes, respectively).

